# Differential impacts of the head on *Platynereis dumerilii* peripheral circadian rhythms

**DOI:** 10.1101/593772

**Authors:** Enrique Arboleda, Martin Zurl, Kristin Tessmar-Raible

## Abstract

**Background:** The marine bristle worm *Platynereis dumerilii* is a useful functional model system for the study of the circadian clock and its interplay with others, e.g. circalunar clocks. The focus has so far been on the worm’s head. However, behavioral and physiological cycles in other animals typically arise from the coordination of circadian clocks located in the brain and in peripheral tissues. Here we focus on peripheral circadian rhythms and clocks, revisit and expand classical circadian work on the worm’s chromatophores, investigate locomotion as read-out and include molecular analyses.

**Results:** We establish that different pieces of the trunk exhibit synchronized, robust oscillations of core circadian clock genes. These circadian core clock transcripts are under strong control of the light-dark cycle, quickly losing synchronized oscillation under constant darkness, irrespective of the absence or presence of heads. Different wavelengths are differently effective in controlling the peripheral molecular synchronization. We have previously shown that locomotor activity is under circadian clock control. Here we show that upon decapitation it still follows the light-dark cycle, but does not free-run under constant darkness. We also observe the rhythmicity of pigments in the worm’s individual chromatophores, confirming that chromatophore size changes follow a circadian pattern. These size changes continue under constant darkness, but cannot be re-entrained by light upon decapitation.

**Conclusions:** Here we provide the first basic characterization of the peripheral circadian clock of *Platynereis dumerilii*. In the absence of the head, light is essential as a major synchronization cue for peripheral molecular and locomotor circadian rhythms. Circadian changes in chromatophore size can however continue for several days in the absence of light/dark changes and the head. Thus, the dependence on the head depends on the type of peripheral rhythm studied. These data show that peripheral circadian rhythms and clocks should be considered when investigating the interactions of clocks with different period lengths, a notion likely also true for other organisms with circadian and non-circadian clocks.

## Introduction

Extensive research focusing on drosophilids and mice showed that the daily behavioral, physiological and metabolic cycles in animals arise from coordination of central circadian clocks located in the brain and peripheral clocks present in multiple tissues (1–3). In Drosophila, several peripheral tissues and appendages (e.g. Malpighian tubules, fat bodies and antennae) have autonomous peripheral clocks that are directly entrained by environmental cycles independent of the central clock, while others, such as oenocytes, are regulated by the circadian clock located in the brain (4,5). The mammalian circadian system is highly hierarchically organized. The master central clock in the suprachiasmatic nucleus (SCN) of the brain (often referred to as a “conductor”) synchronizes internal clock timing to the environmental solar day by passing the information to the peripheral clocks via endocrine and systemic cues (6,7). These peripheral clocks also have self-sustained circadian oscillators, with the master clock coordinating their phase to prevent desynchronization among peripheral tissues, rather than acting as a pacemaker responsible for the periodicity of the cycling itself (8). Besides being phase-controlled by the “SCN conductor”, several mammalian peripheral clocks (e.g. in liver and kidney) have been shown to directly respond to non-photic entrainment cues, like food or exercise (9).

For marine organisms, biorhythms of various period lengths, including circadian and circalunar, have been described across phyla, e.g. as changes in activity levels, coloration and reproductive cycles (reviewed in 10–13). Where studied in detail, like in the marine bristle worm *Platynereis dumerilii* and the marine midge *Clunio marinus*, the light of sun and moon are known to serve as major entrainment cues (14–16). Over recent years, marine rhythms and their possible underlying clockworks have been receiving increasing attention, as the interplay of clocks and rhythms of different organisms is a crucial aspect for ecology (17).

Molecular data on rhythms and clocks in marine invertebrates have become increasingly available over the last decade, now including the bivalves *Mytilus californianus* (18) and *Crassostrea gigas* (19), the sea slugs *Hermissenda crassicornis, Melibe leonina* and *Tritonia diomedea* (20,21), the isopod *Eurydice pulchra* (22–24), the amphipod *Talitrus saltator* (25), the lobsters *Nephrops norvegicus* (26) and *Homarus americanus* (27), the mangrove cricket *Apteronemobius asahinai* (28), the copepods *Calanus finmarchicus* (29) and *Tigriopus californicus* (30), the Antarctic krill *Euphausia superba* (31–34), the Northern krill *Meganyctiphanes norvegica*(27), the marine *midge Clunio marinus* (15,35), and the marine polychaete *Platynereis dumerilii* (14,36). On the marine vertebrate side, especially teleost fish species have been investigated (37–45).

While most of the above mentioned species are difficult to maintain in the laboratory and to investigate at the level of molecular genetics, *P. dumerilii* is a particularly well-established laboratory model (46,47) for marine chronobiological research. It possesses interacting circadian and circalunar clocks and, complementing the molecular work, a detailed analysis of its circadian locomotor activities has been described for adult stages (14,16,48). Evidence of circadian activity also exists for young larval stages within the first days of their development (49). Similar to the isopod *E. pulchra* (22,23), *P. dumerilii* also exhibits a circadian rhythm in its body pigmentation (50–52). This rhythm in pigment cell extension versus contraction was described as a segment-autonomous process (50–52), indicating the presence of autonomous peripheral circadian oscillators.

As *P. dumerilii* beheaded individuals survive well for up to two weeks (53), this feature can be used to study living animals in the absence of its circadian brain clocks. Moreover, *P. dumerilii* has primitive morphological and genetic features, and is hence viewed as evolutionarily slowly evolving (54), a feature which is particularly interesting for understanding the ancestral features of different clocks and rhythms, as well as in the light of the-certainly debated hypothesis-that vertebrates originated from a polychaete-like animal (11).

The work presented here is the first detailed characterization of *P. dumerilii* peripheral circadian rhythms and clocks, covering analyses of transcript level changes of core circadian clock genes, as well as body pigmentation and locomotor activity.

## Results

### *Platynereis dumerilii* peripheral circadian clock gene transcripts quickly desynchronize under complete darkness

As a starting point, we aimed to test for the presence of peripheral clocks in the body of *P. dumerilii* adults. Based on the previously described and tested molecular components of the core circadian oscillator in *P. dumerilii* heads (14), we analyzed the daily transcriptional fluctuation of *bmal, period* and *tr-cry* as representative components of the circadian clock by RT-qPCR (Fig. 1). Daily fluctuation of *bmal, period, tr-cry, timeless, clock and pdp1* transcripts in the trunk are overall similar to those in the head (Fig. 1A,B, for *timeless, clock and pdp1* see Additional file 1: Figure S1 A-C, compare also with (14)). Results were similar irrespective of the segment positions used for analyses, *i.e*. whole trunk (Fig. 1), last 5-7 segments of the body or the adjacent 5-7 segments towards the anterior part of the animal (Additional file 1: Figure S1 D-F) produced consistent results. *Pdu-tr-cry* appears to be the gene that deviates most in this comparison. We attribute this difference to a higher variability of the transcript level synchronization in the trunk given that in most cases it corresponds to the oscillations observed in the head (compare Fig. 1, 2A, 3C, Additional file 1: Figure S1 A-C, and (14)).

**Fig. 1.**
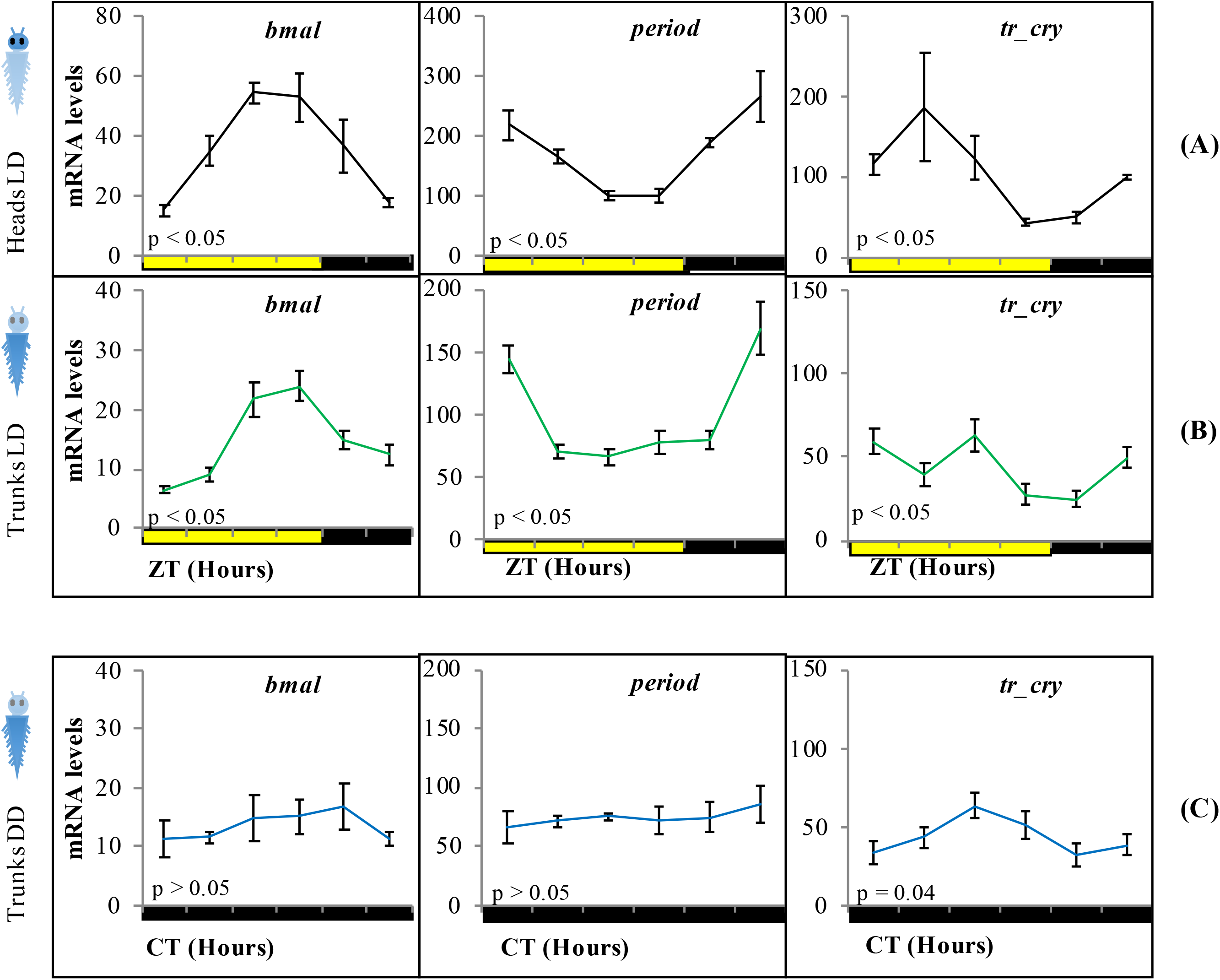
Relative transcript levels of *bmal, period* and *tr-cry* in (A) heads under LD conditions, (B) trunks under LD conditions and (C) trunks in DD conditions on intact animals (*i.e*. not decapitated). ZT = Zeitgeber Time and CT = Circadian Time. p-value estimated on a single factor ANOVA with n=6, 15 and 11 for A, B and C respectively (alpha = 0.05). Error bars denote SEM.

**Fig. 2.**
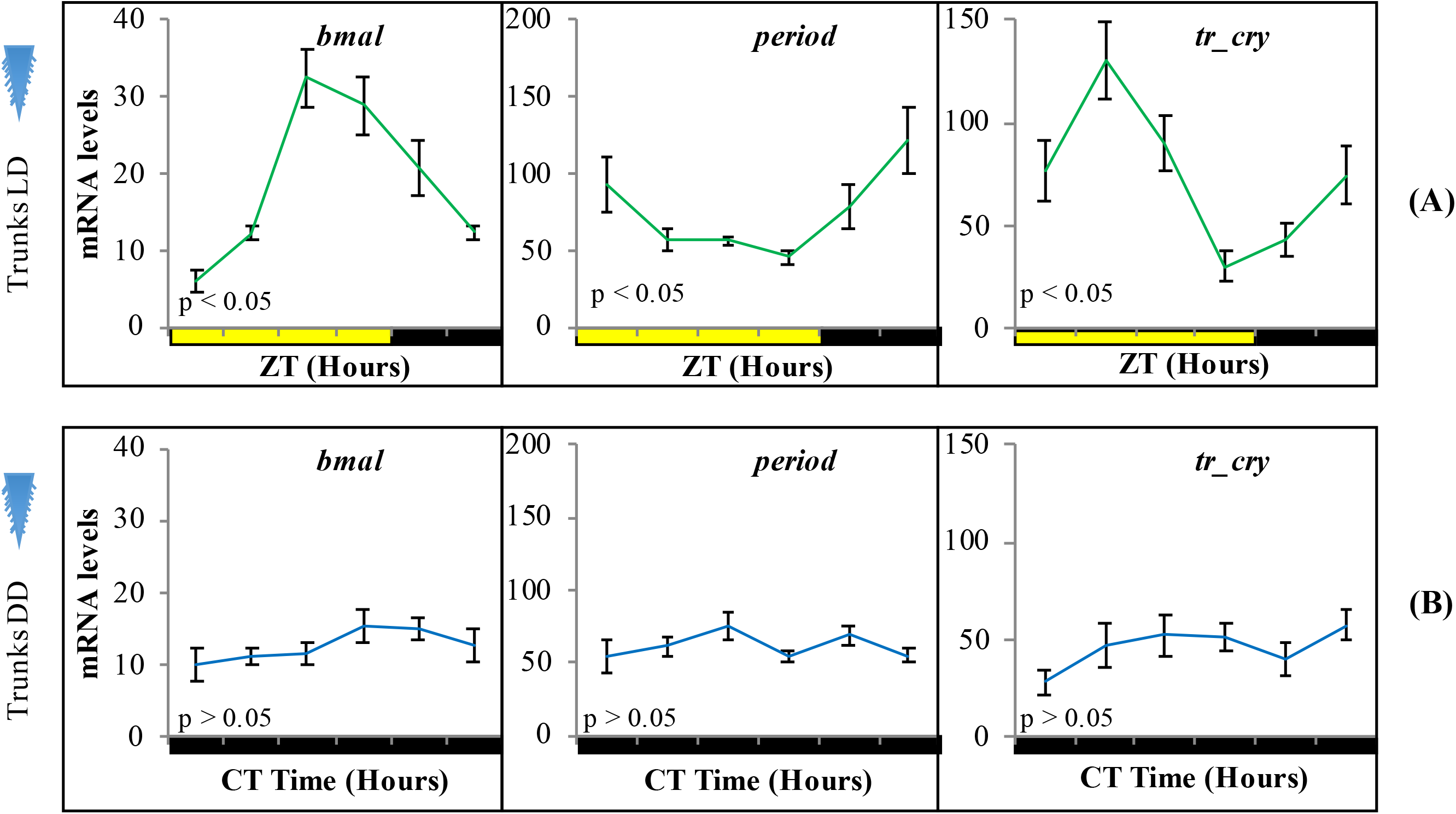
Relative transcript levels of (A) *bmal*, (B) *period* (C) and *tr-cry* in trunks of decapitated animals placed under (A) LD and (B) DD conditions for three days. ZT= Zeitgeber Time. CT= Circadian Time. p-value estimated on a single factor ANOVA with n=6 on each treatment (alpha= 0.05). Error bars denote SEM.

**Fig. 3.**
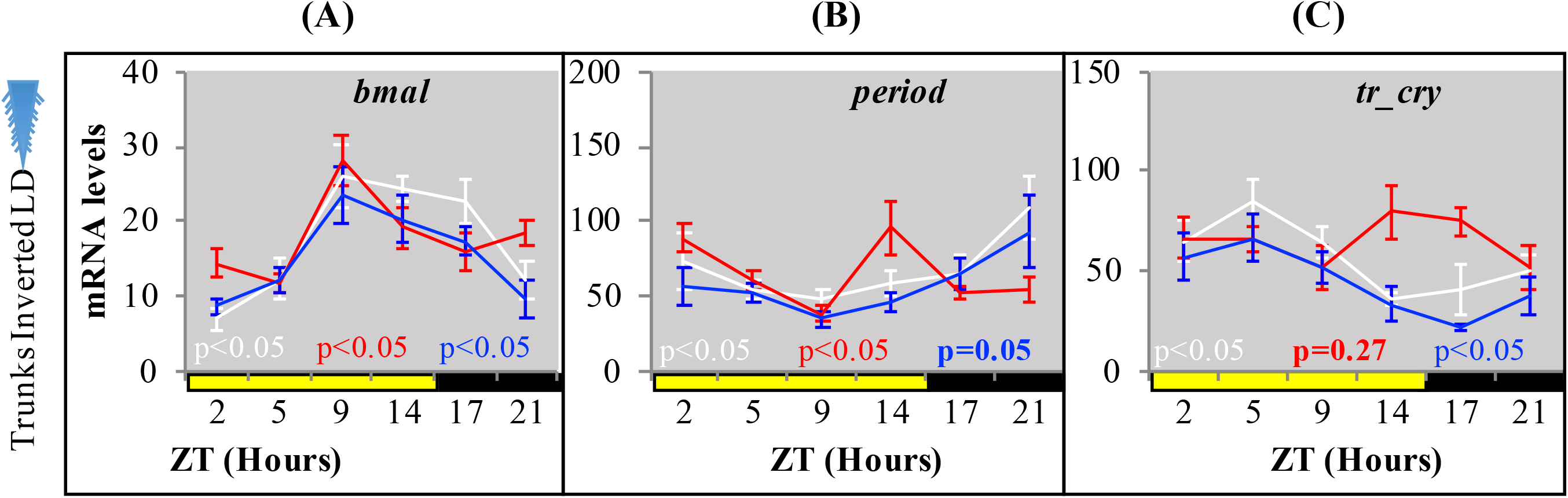
Relative transcript levels of *bmal, period* and *tr-cry* in trunks of decapitated animals placed under inverted LD conditions for seven days with either white, blue or red color (for light spectra and intensity see Additional file 2: Figure S2). ZT= Zeitgeber Time. CT= Circadian Time. p-value estimated on a single factor ANOVA with (alpha= 0.05). Error bars denote SEM.

We next tested if the peripheral circadian clock transcript oscillations would continue synchronously under constant darkness for three days. All tested transcripts dampened strongly, with *tr-cry* still exhibiting weak oscillations (Fig. 1C).

### Light signals are sufficient to maintain circadian clock transcripts in the trunk, independent of the head

To analyze if the peripheral circadian clock transcript oscillations in the worm’s body were dependent on the circadian clock in its head, we took advantage of the fact that bodies of decapitated animals survive in seawater for two weeks (53). Adult animals were decapitated and placed under standard light-dark cycles for three days before RNA extraction. Relative transcript levels of *bmal, period* and *tr-cry* exhibited overall similar levels and continued a clear diel cycling of expression in trunks of decapitated animals (Fig. 2A), indicating that the peripheral circadian clock gene expression continues to synchronously run even in the absence of the brain circadian clock. When we tested core circadian clock transcript oscillations in trunks of decapitated animals under three days of constant darkness, no significant oscillations were detectable (Fig. 2B).

By placing decapitated animals on an inverted light cycle, we next tested for the capacity of peripheral clocks to be re-synchronized by light, again using the transcript oscillations of *bmal, per* and *tr-cry* as readout. Cycling of *bmal* and *per* was re-entrained to the inverted cycle when exposed to white, red or blue light (Fig. 3A,B, for spectra and intensity see in Additional file 2: Figure S2). *tr-cry* transcript oscillations differed from this, in that white and blue light could re-entrain its peripheral oscillations as in the case of *bmal* and *per*, whereas red light did not (Fig. 3C). Overall, these results demonstrate that the peripheral circadian clock transcripts directly respond to changes in the light cycle, independent of the head.

### Chromatophore size follows a circadian pattern and free-runs under constant darkness

In order to assess how our findings on core circadian clock transcripts oscillations might relate to physiology and behavior, we investigated possible outputs of peripheral circadian clocks, starting with changes in chromatophore size in the trunk. Chromatophores are located along the dorsal part of the segmented body of *P. dumerilii*. Based on light microscopy analyses it had previously been shown that the worm’s chromatophores exhibit a segment-autonomous, diel contraction-expansion rhythm with increasing size during the day and decreasing size at night (50–52).

We first aimed at identifying a possibility to automatize the recording of chromatophore changes. We found that chromatophores exhibit a well-detectable autofluorescence under 488 nm light (Fig. 4A,B), which can be used for automatic detection by any image software. In order to characterize the pattern of contraction/expansion of the chromatophores, animals were photographed every three hours over the course of 24h using a fluorescence microscope. We found a clear circadian pattern with higher chromatophore expansion between ZT2 and ZT11 (Fig. 4C), and a sharp drop on chromatophore size from ZT11 to ZT14, before lights go off and the subjective night period starts, already suggestive of an autonomous clock-driven process and not a direct light response, again consistent with previous observations (55).

**Fig. 4.**
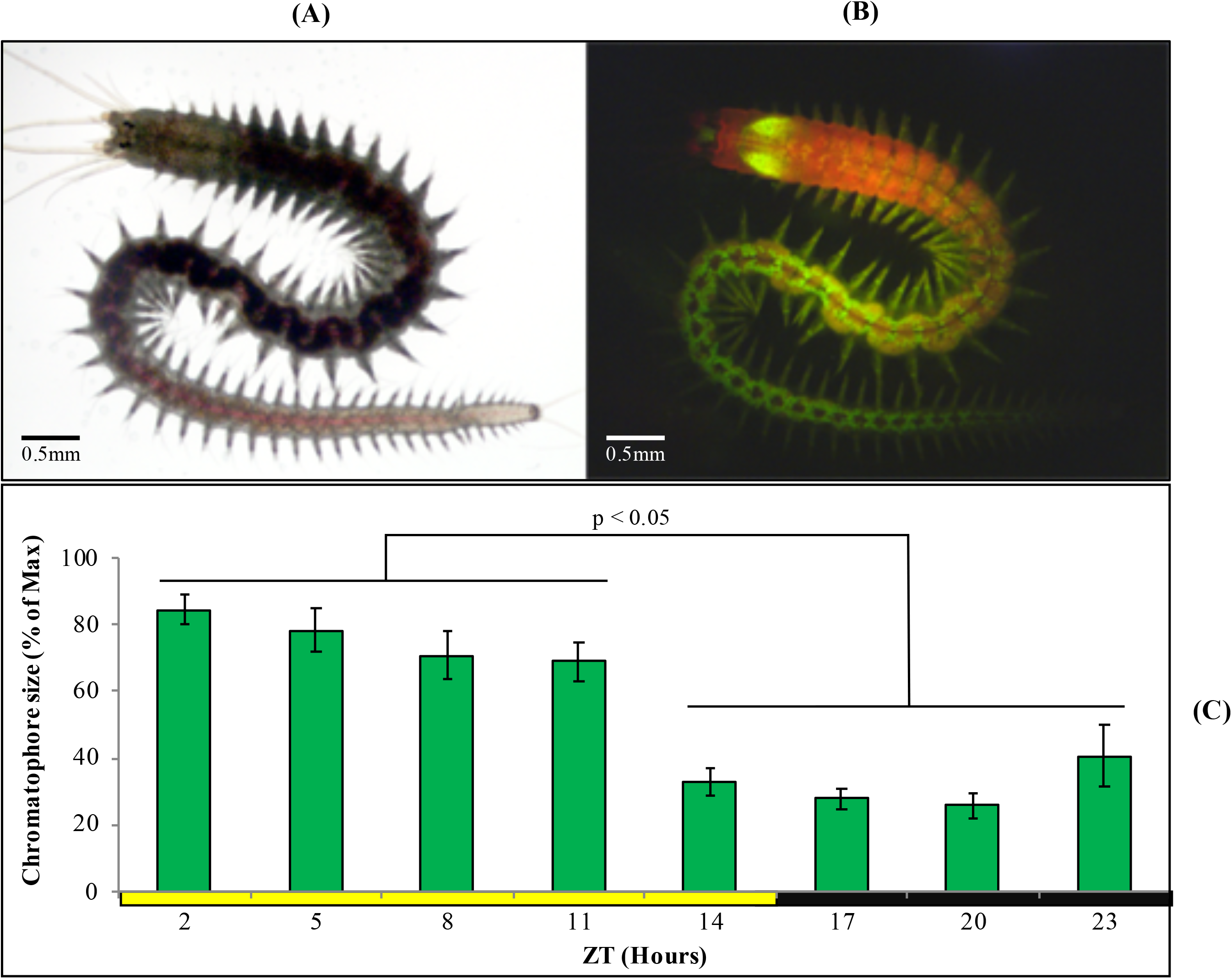
(A) Sexually immature *P. dumerilii* adult under standard light microscopy. (B) Same individual as in (A), now autofluorescent under 488 nm light and a FITC filter, with chromatophores clearly visible as bright green circles (prominent green autofluorescence of the jaws can also be see in the anterior end). (C) Average chromatophore size based on autofluorescence (see materials and methods) of the same group of individuals followed over a 24h period (n=15) under standard LD conditions. Pairwise student’s t-Tests (preceded by F-Tests) were performed comparing each of the ZT hours. Individual averages from ZT2 to ZT5 and ZT14 to ZT23 were statistically similar within them but not between them in all permutations possible (alpha=0.05). Error bars denote SEM.

We next focused on sampling points corresponding to ZT/CT2 and ZT/CT14, during which a ~60% drop in chromatophore size was evident (Fig. 4C, Fig. 5A,B), and used these two time points as a reference for the study of circadian cycling of pigmentation over multiple days. Chromatophores expansion/contraction continued to cycle in animals placed under constant darkness for five consecutive days (Fig. 5C)

For evidence that the light used for measuring chromophore size is not causing re-entrainment in this case, see below.

**Fig. 5.**
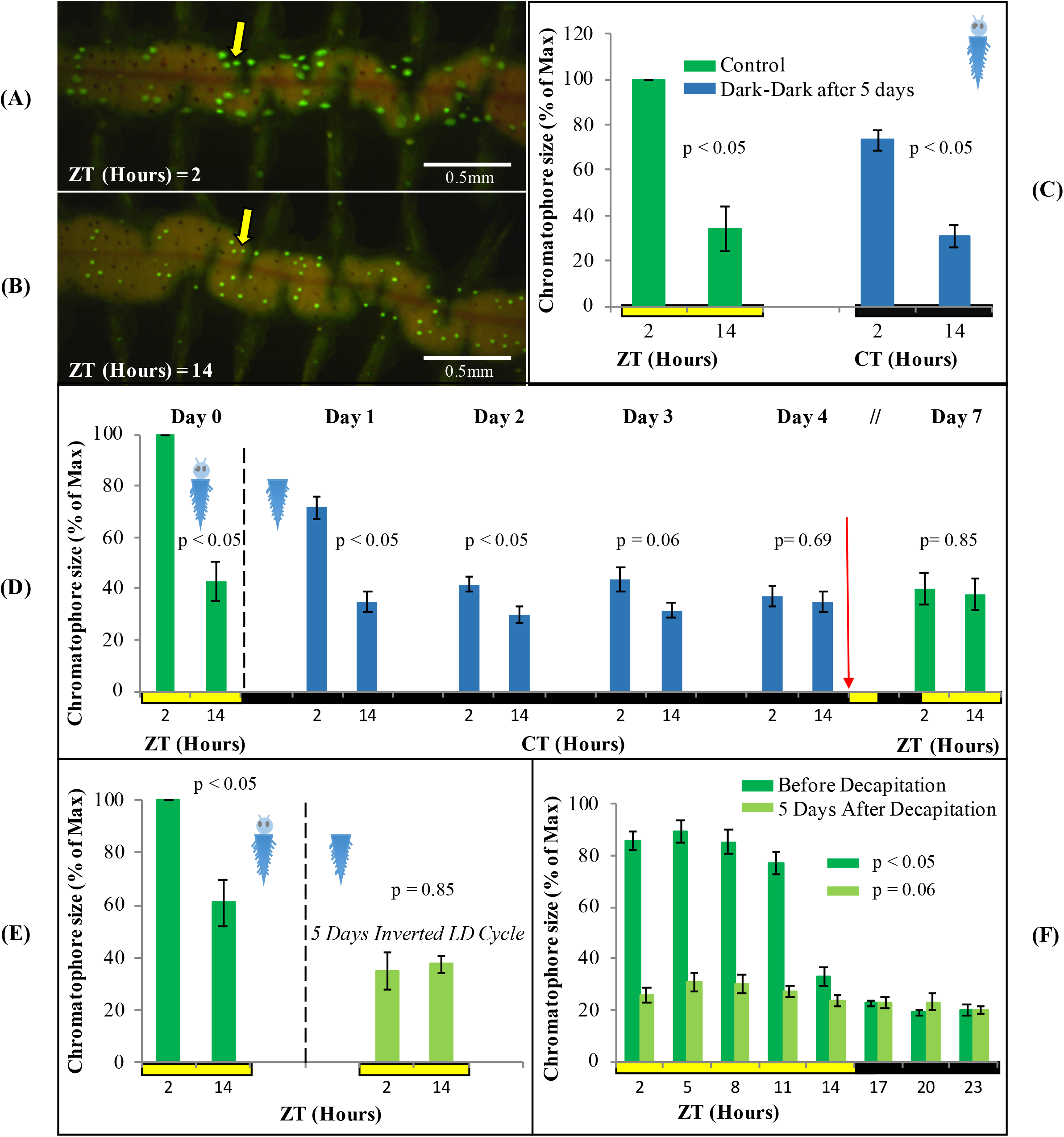
Fluorescent microscopy images of chromatophore size difference between (A) ZT12 and (B) ZT14 under standard LD conditions. (C) Average chromatophore size at ZT/CT 12 and ZT/CT 14 before and after five days under DD conditions (n=4). (D) Average chromatophore size at ZT/CT 12 and ZT/CT 14 over seven consecutive days. Dashed line indicates decapitation and placement in DD conditions. Red arrow indicates re-placement under LD conditions (n=6) (see Additional file 3: Figure S3 for individual replicas). (E) Average chromatophore size, over a 24h period, of individuals before and five days after decapitation (n=10). (F) Average chromatophore size at ZT12 and ZT14 of individuals before and five days after decapitation and placement on an inverted LD cycle (n=6). For (C)(D)(F), pairwise student’s t-Tests (preceded by F-Tests) were performed comparing ZT/CT 2 and ZT/CT 14 at each sampling day. For (E), p-value was estimated on a single factor ANOVA. Error bars denote SEM and alpha= 0.05 in all cases.

### Circadian pattern of chromatophore size free-runs, but cannot be re-entrained by light in decapitated animals

In order to assess if the regulation of the cycling on chromatophore size is governed by peripheral clocks, decapitated animals were used. The same animals were photographed at ZT2 and ZT14 prior to decapitation as a starting reference point. Following decapitation, worms were placed in constant darkness for four days, and subsequently exposed again to a normal LD cycle for additional three days. Chromatophores sizes were registered along the experiment from day 0 to day 4, and again on day 7 to test for possible re-entrainment. Upon decapitation, animals under DD conditions initially continue to exhibit clear rhythms of chromatophore size changes (Fig. 5D, for individual replicas see Additional file 3: Figure S3). Starting with the second day in DD, the rhythm will however start to dampen and become statistically non-significant by day 3 (Fig.5D). Subsequent exposure to a normal LD cycle did not lead to a re-synchronization of the chromatophore rhythm (Fig. 5D). Consistently, cycling of chromatophore size does not get re-entrained on decapitated animals under an inverted LD cycle applied for 5 days (Fig. 5E).

In order to rule out that we may have missed phase-shifts on decapitated animals due to the exposure to the 488nm light during the measurement procedure (due to too low sampling frequency), we also performed a more densely spaced 24-hour sampling on day 5 in LD (post decapitation). This experiment confirmed our interpretation of a dampening of the chromatophore size rhythm and inability to re-entrain in the absence of the head (Fig. 5F). All together, these results suggest a circadian pattern of chromatophore size governed by a peripheral clock, which however requires the head to maintain extended synchronization and for re-entrainment by light.

### Circadian locomotor activity follows the light-dark cycle, but does not free-run under constant darkness in decapitated worms

We next turned to locomotor activity as a read-out for circadian clock activity. We have previously shown that *P. dumerilii* during new moon (circalunar phase of its circalunar clock) exhibits nocturnal circadian locomotor activity, which free-runs under constant darkness for at least three days (14,48). We meanwhile established an automated worm locomotor behavioral tracking system, which measures worm activity as distance moved of the worm’s center point within 6 min time bins (56–58). It should be noted that this new type of analysis measures relative distance moved, compared to the binary (movement: yes or no) manual scoring done by Zantke et al. (14). Intact worms showed a significant circadian rhythmic locomotor activity under LD and DD conditions (Fig. 6A,B, for individual actograms see Additional file 4: Figure S4). In contrast decapitated worms exhibit an overall severe reduction in movement and rhythmicity (Fig.6C,D, for individual actograms see Additional file 5: Figure S5). No significant circadian rhythmicity can be observed under constant darkness in headless worms (Fig.6C,D), with one exception. Overall, these data suggest that lack of signals from the brain lead to a general suppression of movement and lack of circadian information for the locomotor output. Headless worms are not generally paralyzed, as they can show spontaneous bursts of movement (Additional file 5: Figure S5) and- depending on their stage at decapitation-can proceed to maturation and the associated behavioral changes (59). Similar to the rhythmic transcript oscillations of the core circadian clock genes in the trunk, acute light functions as a synchronization cue to the periphery of headless worms, but without head or light stimuli a circadian locomotor pattern cannot be maintained by the trunk alone in 97% of the worms analyzed.

**Fig. 6.**
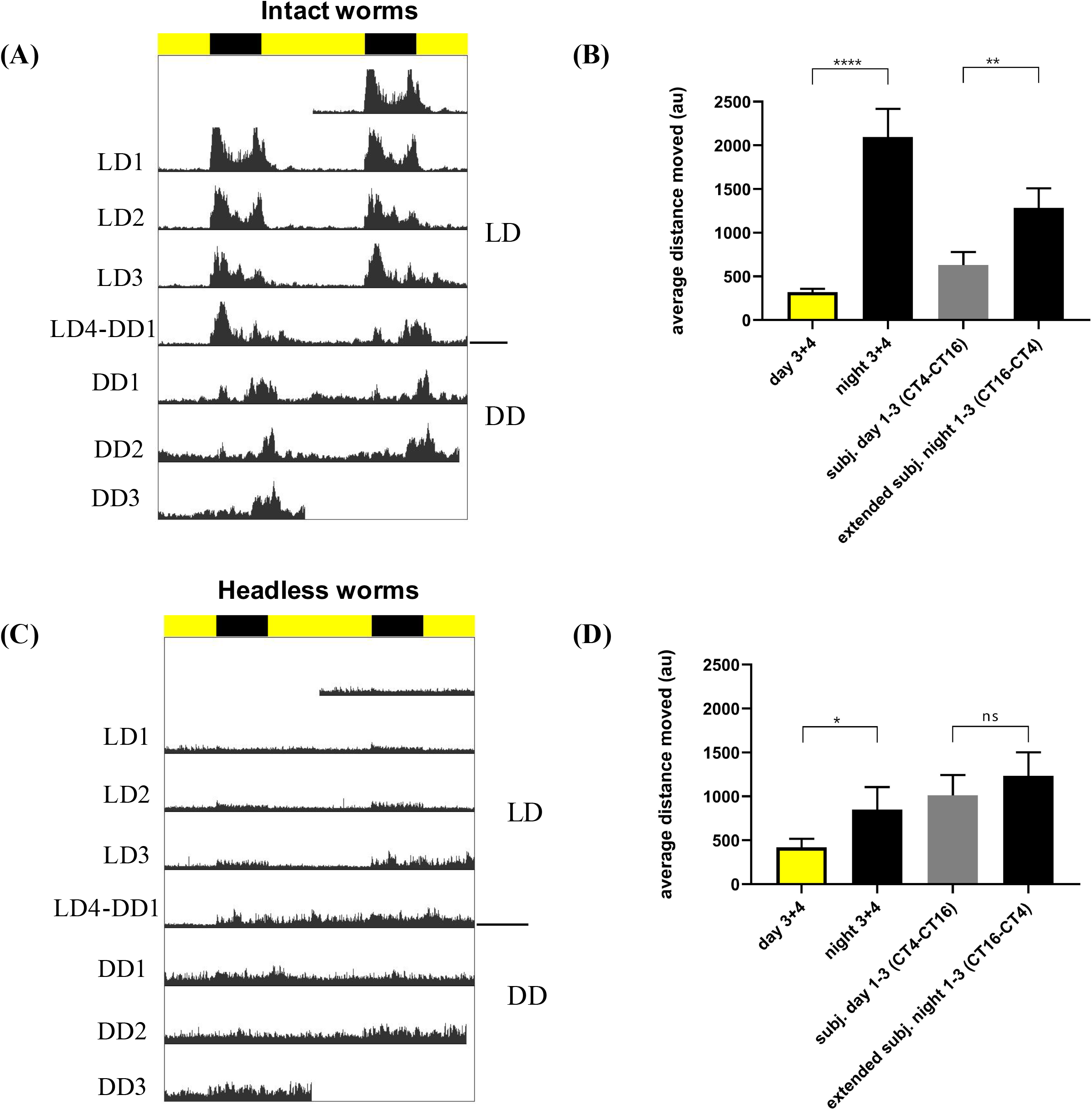
Individual locomotor activity of intact (n=20) and headless (n=30) worms under 16h light and 8h dark (LD) conditions followed by three days of constant darkness (DD). (A,C) Average double plotted actograms of intact (A) and headless (C) worms. Black bar indicates night hours whereas yellow bar indicates day hours during LD conditions. Decapitation of headless worms was performed at ZT14 one day before recording (=LD0). (B,D) Average distance moved of intact (B) and headless worms (D) during day and nighttime, as well as during different time periods of constant darkness (i.e. CT4-CT16 for subjective day, and CT16-CT4 for an extended subjective night). The subjective night period (CT16-CT0) was extended to CT4 because under DD conditions activity levels cycle with a ca. 25h ± 0.22 period (mean ± SEM, see Additional file 4: Figure S4 B), and therefore activity during DD1-3 darkness is expected to run into the early phase of the subjective day. For day and night time activity, only data from LD3 and and LD4 were pooled, because during the first two days after head removal worms hardly moved. Bars indicate mean ± SEM. p-values were calculated using repeated measures ANOVA followed by Sidak’s multiple comparison test with ****p < 0.0001, **p<0.01, *p<0.05

## Discussion

### Molecular peripheral clock

Here we examine peripheral circadian clock transcript changes and diel rhythms in chromatophore size and locomotor behavior in *P. dumerilii* in the absence and presence of its head. An endogenous circadian rhythm driving body pigmentation change in *P. dumerilii* had been previously proposed based on photographic recordings of isolated groups of body segments during the middle of the day and the night (51,52). With the exception of gills in oysters (60), there is no information on fluctuation of circadian clock genes on peripheral tissues or appendages in marine organisms. We document the expression of core circadian clock genes in the periphery of *P. dumerilii*, arguing in favor of a functional circadian peripheral clock and opening an avenue to study the molecular mechanisms of peripheral clocks in marine invertebrates. We show that light/dark cycles can (at least in part) substitute for the head as major synchronizer for continuous peripheral core circadian clock transcript oscillations. We confirm previous work on chromatophore rhythms, which - in contrast to locomotion and transcript oscillations-exhibit free-running rhythms in trunks of headless worms.

As previously reported for the central clock (14), *period* and *bmal* transcripts cycle in antiphase from each other, while *tr-cry* transcripts are neither directly in-phase nor in anti-phase with any of them. Our results on trunks show that cycling of *bmal* and *period* continues under LD conditions, but not under DD independently of the head being present or absent, and can be entrained to an inverted LD cycle on decapitated animals. These results suggest that the peripheral clocks are light dependent (through a yet to be identified set of photoreceptors in the trunk) and get out of synchrony in the different peripheral cells and tissues on the transcriptional level (at least as fast as three days in DD) in the absence of the head. A system with peripheral clocks independent from the central circadian clocks in the brain and entrained directly by environmental signals is reminiscent of *Drosophila melanogaster* (5). In that sense the peripheral clock in *P. dumerilii*resembles that of insects, and light reception in the trunk likely occurs via Cryptochromes and/or Opsins. Candidates include Go-Opsin1 and rOpsin1 (56,61).

Light exhibits different effects on the different readouts. In the cases of transcript oscillations and locomotor activity the head is not required for its impact, suggesting that peripheral photoreceptors mediate this signal. Interestingly, different wavelengths appear to have differential peripheral effects on transcript oscillations (red light being able to re-entrain *per* and *bmal* trunk oscillations, but not *tr-cry)*, which already indicates the involvement of more than one photoreceptor. It will be interesting for future studies to understand why *tr-cry* transcripts behave differently from *bmal* and *per* transcripts under different light conditions. These differences could be the result of *tr-cry* only being expressed in a subset of tissues/cell types that do not desynchronize as quickly.

As in the case of insects (5) and mammals (62), questions regarding how peripheral clocks are entrained and the actual mechanisms that peripheral clock use to drive transcriptional changes on various tissues are still questions to be answered in *P. dumerilii*, as is the case for the specific functions of the peripheral clocks.

It should be noted that our experiments were performed on the complete trunk (or at least several segments), which also leaves open questions regarding the peripheral circadian cycling and their synchronizations in specific segmentally repeated organs and tissues.

### Chromatophore size and the peripheral clock

Daily changes in chromatophore size on *P. dumerilii* can be easily used as a visual read out of the circadian clock, adding up to its locomotor activity, circalunar spawning and clock-related genes as means to study chronobiology in this model system.

Remarkably, the first studies on the cyclic changes of body pigmentation in *P. dumerilii* date back to 1939 (55). Based on this and further classical work, these changes in body pigmentation were already attributed to an endogenous circadian rhythm present in each segment of the worm’s body (50–52), pointing at the existence of peripheral circadian clocks long before their molecular mechanism had been unraveled and circadian peripheral oscillations were proven to exist in the peripheral tissues in mammals (63–66). Our analyses support the classical studies on Platynereis, in that chromatophore size in the body of *P. dumerilii* is higher during day time, with a major drop before the night begins; which argues in favor of a clock-driven manifestation and not a direct light response. The average magnitude of this change corresponds with previous quantitative reports by Fischer (50). We report that individuals placed in DD for five days still exhibit a significant daily difference on chromatophore size, although its maximum value decreases by 25% compared to the initial LD conditions. It has been reported that such cycling starts to fade on individual chromatophores after seven days in DD (50), but we did not test for the long term stability of the cycle for individual chromatophores. Noticeably, a similar circadian cycling of chromatophore size on DD and inverted LD cycles has been shown for the marine isopod *Eurydice pulchra* (23), posing the questions if this regulation is similar.

Changes in chromatophore size with a circadian cycling is commonly seen in crustaceans (23,67–69). There is usually an increase in size during the day thought to be related to UV protection (70), but an inverted pattern of bigger chromatophore size during the night, to possibly enable individuals to camouflage with the substrate, can also be found (71). The fact that *P. dumerilii* is mostly active during the night (14), when pigmentation is lower, does not argue in favor of a camouflage role; especially since pigmentation does not respond to changes in background brightness (50). The most parsimonious option is therefore a role of pigmentation in protecting the animal from UV light. It should be noted, however, that it has also been often proposed, but not tested, that circadian changes in pigmentation might be a mechanism related to energy saving (72).

Remarkably, in our hands the re-entrainment of chromatophore rhythms by light requires the presence of the head. This might be either because the required photoreceptor(s) are located in the head or because hormonal feedback signals, such as Pdu-PDF (pigment dispersing factor) (73), are required for the synchronization process. It is likely that pigmentation in *P. dumerilii* is controlled by a hormonal process, as in some crustaceans (68,69,74), which in turn is governed by the central clock. The hormonal nature of the cycling on chromatophore size has been further supported by the immediate reaction of chromatophores, present on isolated skin tissue, when coelomic fluid from *P. dumerilii* during a given time point (e.g. day or night) is added (52).

Finally, while we overall confirm previous work on the chromatophore rhythms in the trunk of *P. dumerilii*,there is one clear difference between our findings and that of Röseler (52). Her work shows that chromatophore rhythms can still be re-entrained by white light even in the absence of the prostomium (head), whereas in our hands decapitated animals do not re-entrain their chromatophore rhythm in response to white light. We identified two main reasons that might explain this discrepancy. The materials and methods of her paper do not state the light intensity and spectrum. It is thus possible that this strongly deviates from our conditions. The other difference is the extent of head removal. In our study we removed the head including the jaw piece, whereas Röseler (52) specifies prostomium-removal, which implies that her worms still possessed the jaw and the surrounding tissue. This region possesses multiple neurons and neurosecretory cells, which could be important for proper re-entrainment. Further work will certainly help to disentangle these differences.

## Conclusions

We find that the overall circadian clock transcript oscillations of the trunk are under strong control of the light-dark cycle and do not show synchronized oscillation under constant darkness, irrespective of the absence or presence of heads. In the absence of heads, locomotor activity is also strongly coordinated by the light-dark cycle. In contrast, circadian changes of body pigmentation in the trunk free-run over several days in constant darkness, even in the absence of the head. Jointly these data indicate that autonomous peripheral clocks exist in the trunk of the bristle worm, coordinating for instance pigmentation. However, the synchronization of rhythmic circadian oscillations in other peripheral tissues and their respective output are more strongly coordinated by light than by the circadian oscillator positioned in the head of the worm. Our data build a basis for future analyses of the multiple clocks of the bristle worm, but also suggest that peripheral clocks should be taken into consideration when studying other organisms with circadian and non-circadian oscillators.

## Materials and Methods

### Animal cultures

Animals were maintained on controlled temperature and 16:8 hours light-dark (LD) or dark-dark (DD) cycle as previously described (36). Sampling points are presented either as Zeitgeber Time (ZT) for light-dark conditions (LD) or circadian time (CT) for constant darkness (DD).

### Worm decapitation

Animals were anesthetized by adding a few drops of 1M MgCl2 in the seawater until they stopped moving. They were carefully placed on a microscope slide under a binocular dissecting microscope, decapitated, and transferred to fresh seawater again. Decapitation was done with sterile surgical blades cutting on the first segments below the pharyngeal region in order to collect only the posterior part of the body (*i.e*. the trunk). They were consistently performed between ZT 13.5 and ZT14. For head samples, the region containing the pharynx and the posterior end of the head was removed (see (14)).

### Circadian re-entrainment under white, blue and red light conditions

When testing for circadian re-entrainment (*i.e*. an inverted light cycle) under white, blue or red light, animals were exposed to the new conditions for seven days before sampling. Light spectra and intensities of white, blue and red LEDs (ProfiLuxSimu-L from GHL advanced technology gmbh, Germany) used for circadian re-entrainment were measured using an ILT950 spectrometer (International Light Technologies Inc., Peabody, USA). Special care was taken to account for the standard conditions where the worms were housed, *i.e*. 22cm away from light source and with a transparent plastic lid positioned between the spectrometer and the light source. Measured light intensities for white, red and blue lights were 8,2 × 10^13^ photons/cm^2^/s, 3,8 × 10^13^ photons/cm^2^/s and 2,4 × 10^13^ photons/cm^2^/s, respectively (for spectra see Additional file 2: Figure S2A).

### Total RNA extraction and RT-qPCR

Total RNA was extracted from heads or trunks (i.e. decapitated animals) using the RNeasy Mini Kit (QIAGEN). Reverse transcription was carried out using 0.4 μg of total RNA as template (QuantiTect Reverse Transcription kit, QIAGEN). RT-qPCR analyses were performed using a Step-One-Plus cycler. The expression of each test gene was normalized by the amount of the internal control gene cdc5. The relative expression was calculated using the formula 1/2^ΔCt^. Primers and PCR program used are listed in Zantke et al. (14).

### Chromatophore size

Three consecutive segments located towards the middle of the body were selected on each animal to evaluate changes in chromatophore size. In order to precisely re-identify the same segments over the course of the experiments, animals were anesthetized with MgCl2 and a parapodium, two segments away from the region of interest, was removed with a sterile surgical blade. When required, one animal at a time was placed on a glass cover without water and extended carefully. An epifluorescence stereoscope (Zeiss Lumar) with a 488 nm laser and a FITC filter was used to take pictures making sure to always use the same magnification across animals and sampling points. Animals were placed again in seawater until the next sampling point. Image analysis was done using Adobe Photoshop. On fluorescent images, the RGB channels red and blue were lowered to zero, and the three segments of interest were extracted by erasing the unwanted area (chromatophores on each segment have a specific pattern, which makes their visual identification easier). A new layer was generated by using the magic wand tool to single out the bright green chromatophores from the background fluorescence. Using the image’s histogram, the number of colored pixels was used as a proxy of chromatophore size. Animals had between 20 and 40 chromatophores along the three segments, but no effort was done to quantify size of individual chromatophores. Instead, the sum of all the chromatophores of interest were used as chromatophore size value at each time point. Absolute pixel number was expressed as a percentage of the maximum value for each animal across all time points of the experiment (i.e. x_i_/(Max{x_i_, …, x_j_}*100). Average among biological replica and SEM were further calculated.

### Locomotor Activity Assay

Immature worms of comparable size were starved for three days before the start of the assay. After decapitation, worms were placed in individual hemispherical concave wells (diameter = 35 mm, depth = 15 mm) of a custom-made 36-well clear plastic plate (as described in (56)). Intact worms were also treated with MgCl2 for 5min prior to locomotor recording to ensure proper comparisons to decapitated worms. Video recording of worm’s behavior over several days was accomplished as described previously (14), using an infrared (λ = 990 nm) LED array (Roschwege GmbH) illuminating the behavioral chamber and an infrared high-pass filter restricting the video camera. White light was generated by custom made LEDs (Marine Breeding Systems, St. Gallen, Switzerland) with the intensity set to 5,2 × 10^14^ photons/cm^2^/s at the place were worms were housed (for light spectrum see Additional file 2: Figures S2B). Trajectories of locomotor activity of individual worms were deduced from the video recordings by an automated tracking software developed by LoopBio gmbh (57, 58). Locomotor activity trajectories reflect the distance moved of each worm’s center point across 6min time bins. Activity data was plotted as double-plotted actograms using the ActogramJ plugin for Fiji (75).

### Statistical Analyses

Statistical analyses were performed using either the Data Analysis plug-in in Microsoft Excel using an alpha value of 0.05 (molecular and chromatophore data) or GraphPad Prism (locomotor activity data). For changes in chromatophore size, each two-sample two-tailed student’s t-Test was preceded by an F-Test to check if the variances of the two groups were equal or not. To test if transcriptional changes in gene expression over time oscillated to a statistically significant difference, fold change data was analyzed for each sampling point using single factor ANOVA. In order to ease the logistic process of analyzing a considerable amount of data, RNA samples and chromatophore size images were not analyzed blindly but chronologically as experiments were being performed. Statistical differences in locomotor activity across treatments were estimated using repeated measures ANOVA followed by Sidak’s multiple comparison test. To identify the free-running period length of intact worms under DD conditions Lomb-Scargle periodogram analysis was done using the ActogramJ plugin for Fiji (75).

## Ethics approval and consent to participate

All animal work was conducted according to Austrian and European guidelines for animal research.

## Consent for publication

Not applicable.

## Availability of data and material

All sequence resources referred to here were already published previously and submitted to public databases. All other data that support the findings of this study are available from the corresponding author upon reasonable request.

## Authors’ contributions

EA and KTR designed the study and wrote the manuscript. MZ reviewed and commented on manuscript. EA and MZ performed experiments: MZ – locomotor activity experiments and measurement of light spectra and intensities, EA – all other experiments. EA, KT and MZ did data analysis, interpretation and discussion.

## Acknowledgements

We thank the members of the Tessmar-Raible and Raible groups for discussions, especially Juliane Zantke, Vinoth Babu Veedin Rajan, Birgit Pöhn and Max Hofbauer for sharing details on the automated locomotor behavioral tracking prior to publication. Andreij Belokurov and Margaryta Borysova for excellent worm care at the MFPL aquatic facility and Claudia Lohs and Katharina Schipany for excellent technical assistance.

## Additional Material

File name: Additional file 1 File format: PDF Title of data:

**Figure S1.**
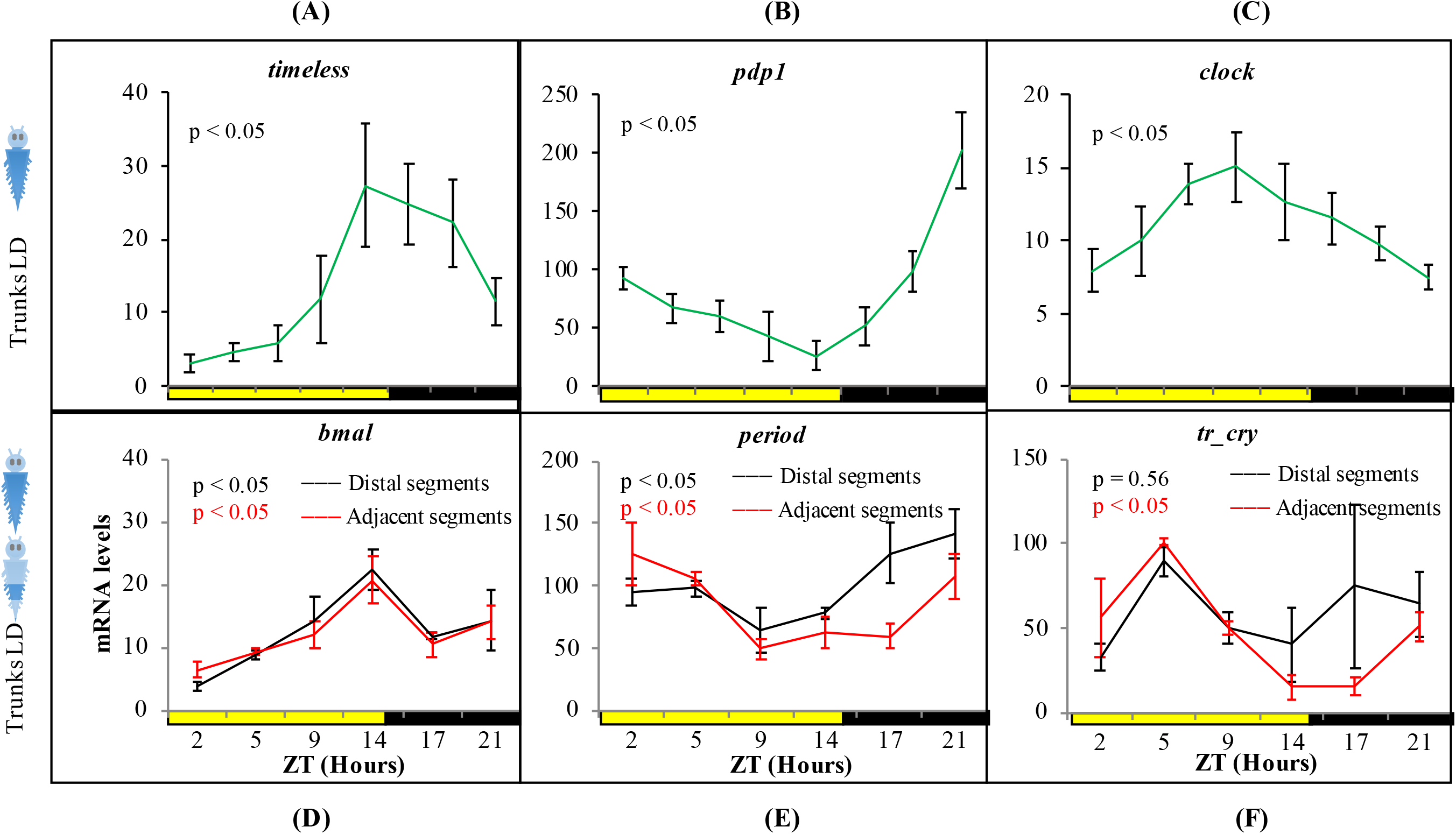
Description of data: Relative transcript levels of (A) *timeless*, (B) *pdp1* and (C) *clock* in trunks of intact animals *(i.e*. not decapitated) under standard LD conditions (n= 5 to 8 per ZT point); and of (D) bmal, (E) period and (F) tr-cry on the last 5-7 segments of the body (black line) or the adjacent 5-7 segments towards the anterior part of the animal (red line) (n=3 in all cases). ZT= Zeitgeber Time. p-value estimated on a single factor ANOVA (alpha= 0.05). Error bars denote SEM.

File name: Additional file 2 File format: PDF Title of data:

**Figure S2.**
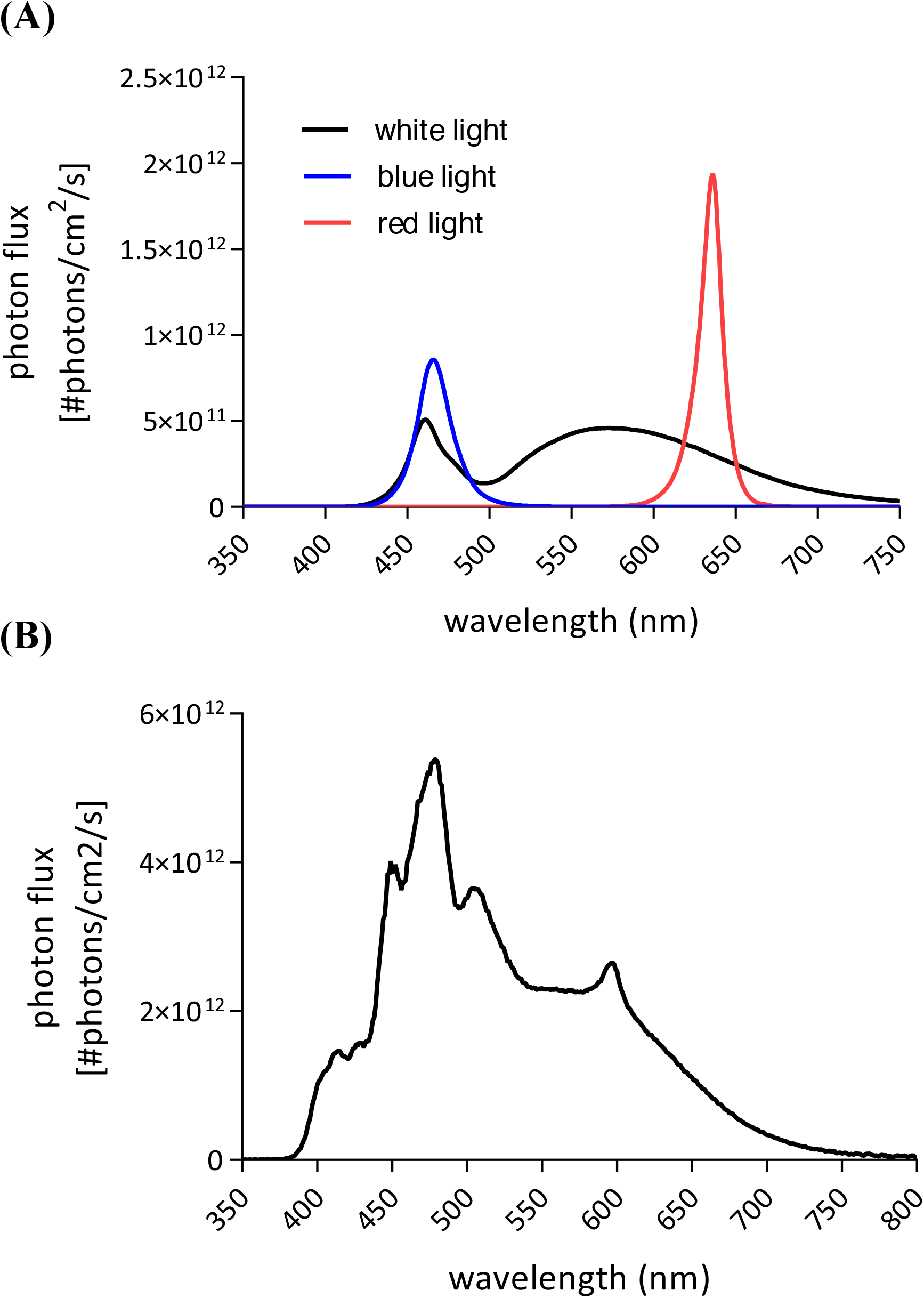
Description of data: (A) Spectra and intensity of light used for re-entrainment of animals (see Figs. 2-3 in the main text). (B) White light spectrum used in the locomotor activity assay.

File name: Additional file 3 File format: PDF Title of data:

**Figure S3.**
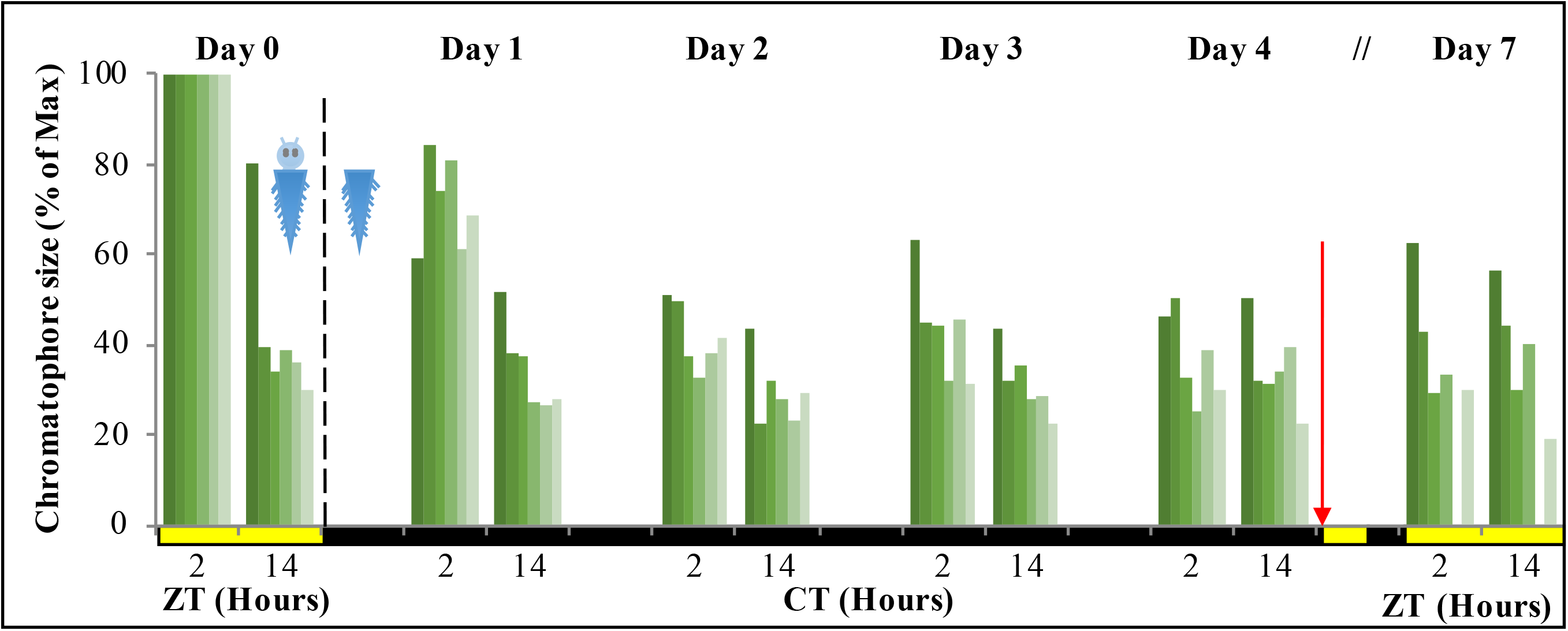
Description of data: Individual replicas of animals used on Fig. 5C, showing chromatophore size at ZT/CT 12 and ZT/CT 14 over seven consecutive days. Dashed line indicates decapitation and placement in DD conditions. Red arrow indicates re-placement under LD conditions (n=6). ZT= Zeitgeber Time and CT= Circadian Time.

File name: Additional file 4 File format: PDF Title of data:

**Figure S4.**
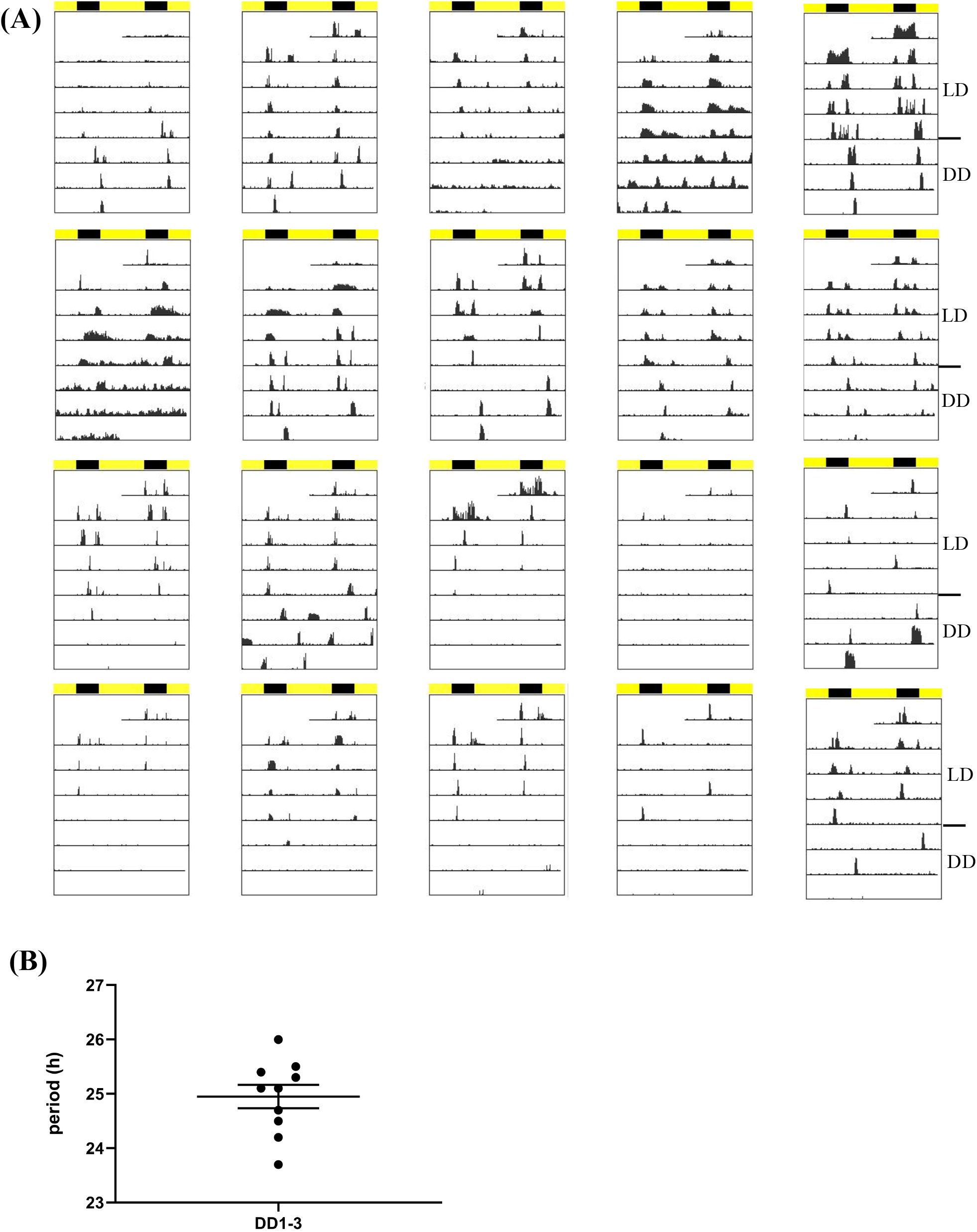
(A) Individual locomotor activity of intact worms under 16h light and 8h dark (LD) conditions followed by three days of constant darkness (DD) depicted in individual double plotted actograms. Black bar indicates night hours whereas yellow bar indicates day hours during LD conditions. Decapitation of headless worms was performed at ZT14 one day before recording (=LD0). (B) Lomb-Scargle periodogram analysis of activity data recorded under DD1-3 reveals a circadian free running period of 25.0h ± 0.22 h (n=10, mean ± SEM). Only periods that had a power value >20 were considered (power values of >10 were already classified to be significantly rhythmic (p<0.05) by the Lomb-Scargle analyses, but we used a more stringent cut-off to reduce the probability of detecting false-positive periods). Out of the 20 worms tested only the first 10 worms depicted in (A) show power values >20, and therefore only these were considered to calculate the mean period length.

File name: Additional file 5 File format: PDF Title of data:

**Figure S5.**
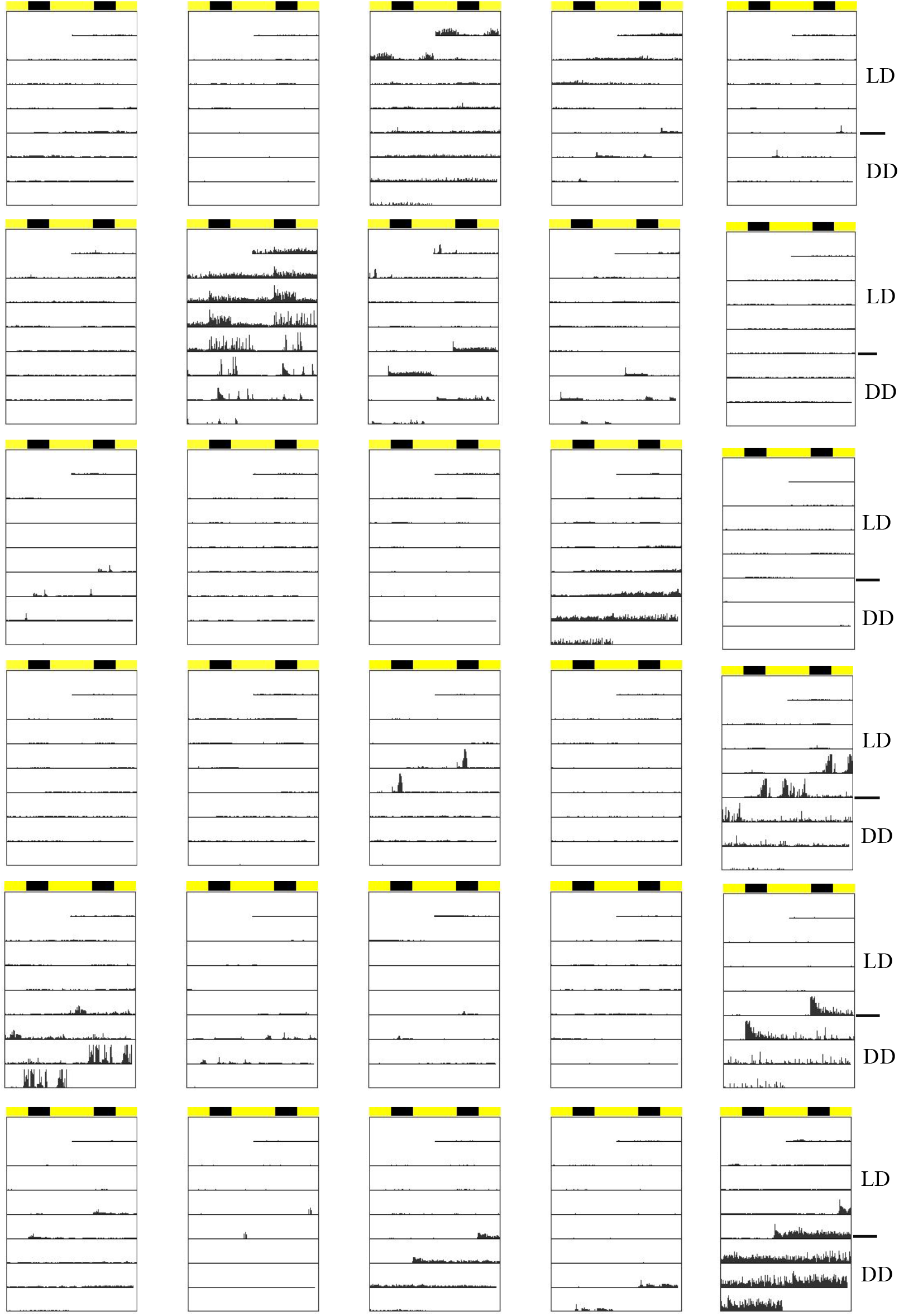
Description of data: Individual locomotor activity of headless worms under 16h light and 8h dark (LD) conditions followed by three days of constant darkness (DD) depicted in individual double plotted actograms. Black bar indicates night hours whereas yellow bar indicates day hours during LD conditions. Decapitation of headless worms was performed at ZT14 one day before recording (=LD0)

## References

1. Mure LS, Le HD, Benegiamo G, Chang MW, Rios L, Jillani N, et al. Diurnal transcriptome atlas of a primate across major neural and peripheral tissues. Science. 2018 16;359(6381).

2. Pilorz V, Helfrich-Förster C, Oster H. The role of the circadian clock system in physiology. Pflugers Arch-Eur J Physiol. 2018 Feb 1;470(2):227–39.

3. Richards J, Gumz ML. Advances in understanding the peripheral circadian clocks. FASEB J. 2012 Sep;26(9):3602–13.

4. Ito C, Tomioka K. Heterogeneity of the Peripheral Circadian Systems in Drosophila melanogaster: A Review. Front Physiol. 2016;7:8.

5. Tomioka K, Uryu O, Kamae Y, Umezaki Y, Yoshii T. Peripheral circadian rhythms and their regulatory mechanism in insects and some other arthropods: a review. J Comp Physiol B, Biochem Syst Environ Physiol. 2012 Aug;182(6):729–40.

6. Mohawk JA, Green CB, Takahashi JS. Central and peripheral circadian clocks in mammals. Annu Rev Neurosci. 2012;35:445–62.

7. Partch CL, Green CB, Takahashi JS. Molecular architecture of the mammalian circadian clock. Trends Cell Biol. 2014 Feb;24(2):90–9.

8. Yoo S-H, Yamazaki S, Lowrey PL, Shimomura K, Ko CH, Buhr ED, et al. PERIOD2::LUCIFERASE real-time reporting of circadian dynamics reveals persistent circadian oscillations in mouse peripheral tissues. Proc Natl Acad Sci U S A. 2004 Apr 13;101(15):5339–46.

9. Tahara Y, Shibata S. Entrainment of the mouse circadian clock: Effects of stress, exercise, and nutrition. Free Radical Biology and Medicine. 2018 May 1;119:129–38.

10. Bulla M, Oudman T, Bijleveld AI, Piersma T, Kyriacou CP. Marine biorhythms: bridging chronobiology and ecology. Philos Trans R Soc Lond, B, Biol Sci. 2017 Nov 19;372(1734).

11. Last KS, Hendrick VJ. The Clock-Work Worms: Diversity and Function of Clock Expression in Marine Polychaete Worms. In: Annual, Lunar, and Tidal Clocks [Internet]. Springer, Tokyo; 2014 [cited 2018 Jun 19]. p. 179–99. Available from: https://link.springer.com/chapter/10.1007/978-4-431-55261-1_10

12. Tessmar-Raible K, Raible F, Arboleda E. Another place, another timer: Marine species and the rhythms of life. Bioessays. 2011 Mar 1;33(3):165–72.

13. Raible F, Takekata H, Tessmar-Raible K. An Overview of Monthly Rhythms and Clocks. Front Neurol [Internet]. 2017 [cited 2018 Dec 31];8.Available from: https://www.frontiersin.org/articles/10.3389/fneur.2017.00189/full

14. Zantke J, Ishikawa-Fujiwara T, Arboleda E, Lohs C, Schipany K, Hallay N, et al. Circadian and Circalunar Clock Interactions in a Marine Annelid. Cell Reports. 2013 Oct 17;5(1):99–113.

15. Kaiser TS, Poehn B, Szkiba D, Preussner M, Sedlazeck FJ, Zrim A, et al. The genomic basis of circadian and circalunar timing adaptations in a midge. Nature. 2016 01;540(7631):69–73.

16. Hauenschild C. Lunar periodicity. Cold Spring Harb Symp Quant Biol. 1960;25:491–7.

17. Schwartz William J., Helm Barbara, Gerkema Menno P. Wild clocks: preface and glossary. Philosophical Transactions of the Royal Society B: Biological Sciences. 2017 Nov 19;372(1734):20170211.

18. Connor KM, Gracey AY. Circadian cycles are the dominant transcriptional rhythm in the intertidal mussel Mytilus californianus. Proc Natl Acad Sci USA. 2011 Sep 20;108(38):16110–5.

19. Perrigault M, Tran D. Identification of the Molecular Clockwork of the Oyster Crassostrea gigas. PLOS ONE. 2017 Jan 10;12(1):e0169790.

20. Cook GM, Gruen AE, Morris J, Pankey MS, Senatore A, Katz PS, et al. Sequences of Circadian Clock Proteins in the Nudibranch Molluscs Hermissenda crassicornis, Melibe leonina, and Tritonia diomedea. Biol Bull. 2018 Jun;234(3):207–18.

21. Duback VE, Sabrina Pankey M, Thomas RI, Huyck TL, Mbarani IM, Bernier KR, et al. Localization and expression of putative circadian clock transcripts in the brain of the nudibranch Melibe leonina. Comp Biochem Physiol, Part A Mol Integr Physiol. 2018 Sep;223:52–9.

22. Wilcockson DC, Zhang L, Hastings MH, Kyriacou CP, Webster SG. A novel form of pigment-dispersing hormone in the central nervous system of the intertidal marine isopod, Eurydice pulchra (leach). Journal of Comparative Neurology. 2011;519(3):562–75.

23. Zhang L, Hastings MH, Green EW, Tauber E, Sladek M, Webster SG, et al. Dissociation of Circadian and Circatidal Timekeeping in the Marine Crustacean Eurydice pulchra. Curr Biol. 2013 Oct 7;23(19):1863–73.

24. O’Neill JS, Lee KD, Zhang L, Feeney K, Webster SG, Blades MJ, et al. Metabolic molecular markers of the tidal clock in the marine crustacean Eurydice pulchra. Curr Biol. 2015 Apr 20;25(8):R326-327.

25. Hoelters L, O’Grady JF, Webster SG, Wilcockson DC. Characterization, localization and temporal expression of crustacean hyperglycemic hormone (CHH) in the behaviorally rhythmic peracarid crustaceans, Eurydice pulchra (Leach) and Talitrus saltator (Montagu). Gen Comp Endocrinol. 2016 01;237:43–52.

26. Sbragaglia V, Lamanna F, Mat AM, Rotllant G, Joly S, Ketmaier V, et al. Identification, Characterization, and Diel Pattern of Expression of Canonical Clock Genes in Nephrops norvegicus (Crustacea: Decapoda) Eyestalk. PLoS ONE. 2015;10(11):e0141893.

27. Christie AE, Yu A, Roncalli V, Pascual MG, Cieslak MC, Warner AN, et al. Molecular evidence for an intrinsic circadian pacemaker in the cardiac ganglion of the American lobster, Homarus americanus-Is diel cycling of heartbeat frequency controlled by a peripheral clock system? Mar Genomics. 2018 Oct;41:19–30.

28. Takekata H, Matsuura Y, Goto SG, Satoh A, Numata H. RNAi of the circadian clock gene period disrupts the circadian rhythm but not the circatidal rhythm in the mangrove cricket. Biol Lett. 2012 Aug 23;8(4):488–91.

29. Häfker NS, Meyer B, Last KS, Pond DW, Hüppe L, Teschke M. Circadian Clock Involvement in Zooplankton Diel Vertical Migration. Current Biology. 2017 Jul 24;27(14):2194–2201.e3.

30. Nesbit KT, Christie AE. Identification of the molecular components of a Tigriopus californicus (Crustacea, Copepoda) circadian clock. Comp Biochem Physiol Part D Genomics Proteomics. 2014 Dec;12:16–44.

31. Biscontin A, Wallach T, Sales G, Grudziecki A, Janke L, Sartori E, et al. Functional characterization of the circadian clock in the Antarctic krill, Euphausia superba. Scientific Reports. 2017 Dec 18;7(1):17742.

32. Pittà CD, Biscontin A, Albiero A, Sales G, Millino C, Mazzotta GM, et al. The Antarctic Krill Euphausia superba Shows Diurnal Cycles of Transcription under Natural Conditions. PLOS ONE. 2013 Jul 17;8(7):e68652.

33. Teschke M, Wendt S, Kawaguchi S, Kramer A, Meyer B. A circadian clock in Antarctic krill: an endogenous timing system governs metabolic output rhythms in the euphausid species Euphausia superba. PLoS ONE. 2011;6(10):e26090.

34. Mazzotta GM, De Pittà C, Benna C, Tosatto SCE, Lanfranchi G, Bertolucci C, et al. A cry from the krill. Chronobiol Int. 2010 May;27(3):425–45.

35. Kaiser TS, Heckel DG. Genetic Architecture of Local Adaptation in Lunar and Diurnal Emergence Times of the Marine Midge Clunio marinus (Chironomidae, Diptera). PLoS One [Internet]. 2012 Feb 22 [cited 2018 Jul 23];7(2). Available from: https://www.ncbi.nlm.nih.gov/pmc/articles/PMC3285202/

36. Schenk S, Bannister SC, Sedlazeck FJ, Anrather D, Minh BQ, Bileck A, et al. Combined transcriptome and proteome profiling reveals specific molecular brain signatures for sex, maturation and circalunar clock phase. Elife. 2019 Feb 15;8.

37. Okano K, Ozawa S, Sato H, Kodachi S, Ito M, Miyadai T, et al. Light-and circadian-controlled genes respond to a broad light spectrum in Puffer Fish-derived Fugu eye cells. Sci Rep. 2017 18;7:46150.

38. Hur S-P, Takeuchi Y, Esaka Y, Nina W, Park Y-J, Kang H-C, et al. Diurnal expression patterns of neurohypophysial hormone genes in the brain of the threespot wrasse Halichoeres trimaculatus. Comp Biochem Physiol, Part A Mol Integr Physiol. 2011 Apr;158(4):490–7.

39. Mogi M, Uji S, Yokoi H, Suzuki T. Early development of circadian rhythmicity in the suprachiamatic nuclei and pineal gland of teleost, flounder (Paralichthys olivaeus), embryos. Dev Growth Differ. 2015 Aug;57(6):444–52.

40. Park J-G, Park Y-J, Sugama N, Kim S-J, Takemura A. Molecular cloning and daily variations of the Period gene in a reef fish Siganus guttatus. J Comp Physiol A Neuroethol Sens Neural Behav Physiol. 2007 Apr;193(4):403–11.

41. Rhee J-S, Kim B-M, Lee B-Y, Hwang U-K, Lee YS, Lee J-S. Cloning of circadian rhythmic pathway genes and perturbation of oscillation patterns in endocrine disrupting chemicals (EDCs)-exposed mangrove killifish Kryptolebias marmoratus. Comp Biochem Physiol C Toxicol Pharmacol. 2014 Aug;164:11–20.

42. Sánchez JA, Madrid JA, Sánchez-Vázquez FJ. Molecular cloning, tissue distribution, and daily rhythms of expression of per1 gene in European sea bass (Dicentrarchus labrax). Chronobiol Int. 2010 Jan;27(1):19–33.

43. Toda R, Okano K, Takeuchi Y, Yamauchi C, Fukushiro M, Takemura A, et al. Hypothalamic expression and moonlight-independent changes of Cry3 and Per4 implicate their roles in lunar clock oscillators of the lunar-responsive Goldlined spinefoot. PLoS ONE. 2014;9(10):e109119.

44. Vera LM, Negrini P, Zagatti C, Frigato E, Sánchez-Vázquez FJ, Bertolucci C. Light and feeding entrainment of the molecular circadian clock in a marine teleost (Sparus aurata). Chronobiol Int. 2013 Jun;30(5):649–61.

45. Watanabe N, Itoh K, Mogi M, Fujinami Y, Shimizu D, Hashimoto H, et al. Circadian pacemaker in the suprachiasmatic nuclei of teleost fish revealed by rhythmic period2 expression. Gen Comp Endocrinol. 2012 Sep 1;178(2):400–7.

46. Fischer AH, Henrich T, Arendt D. The normal development of Platynereis dumerilii (Nereididae, Annelida). Frontiers in Zoology. 2010 Dec 30;7:31.

47. Fischer A, Dorresteijn A. The polychaete Platynereis dumerilii (Annelida): a laboratory animal with spiralian cleavage, lifelong segment proliferation and a mixed benthic/pelagic life cycle. BioEssays. 2004;26(3):314–25.

48. Zantke J, Oberlerchner H, Tessmar-Raible K. Circadian and Circalunar Clock Interactions and the Impact of Light in Platynereis dumerilii. In: Numata H, Helm B, editors. Annual, Lunar, and Tidal Clocks: Patterns and Mechanisms of Nature’s Enigmatic Rhythms [Internet]. Tokyo: Springer Japan; 2014 [cited 2018 Dec 30]. p. 143–62. Available from: https://doi.org/10.1007/978-4-431-55261-1_8

49. Tosches MA, Bucher D, Vopalensky P, Arendt D. Melatonin Signaling Controls Circadian Swimming Behavior in Marine Zooplankton. Cell. 2014 Sep 25;159(1):46–57.

50. Fischer A. Über die Chromatophoren und den Farbwechsel bei dem Polychäten Platynereis dumerilii. Zeitschrift für Zellforschung und Mikroskopische Anatomie. 1964 Mar 1;65(2):290–312.

51. Röseler I. Untersuchungen über den physiologischen Farbwechsel bei dem Polychaeten Platynereis dumerilii. Zool Anz, Suppl-Bd 33, Verh Zool Ges. 1969;267–73.

52. Röseler I. Die Rhythmik der Chromatophoren des Polychaeten Platynereis dumerilii. Zeitschrift für vergleichende Physiologie. 1970 Jun 1;70(2):144–74.

53. Hofmann DK. Regeneration and endocrinology in the polychaetePlatynereis dumerilii: An experimental and structural study. Wilehm Roux Arch Dev Biol. 1976 Mar;180(1):47–71.

54. Tessmar-Raible K, Arendt D. Emerging systems: between vertebrates and arthropods, the Lophotrochozoa. Current Opinion in Genetics & Development. 2003 Aug 1;13(4):331–40.

55. Hempelmann F. Chromatophoren bei Nereis. Zeitschr Wiss Zool Abt A. 1939;152:353–83.

56. Ayers T, Tsukamoto H, Gühmann M, Veedin Rajan VB, Tessmar-Raible K. A Go-type opsin mediates the shadow reflex in the annelid Platynereis dumerilii. BMC Biol [Internet]. 2018 Apr 18 [cited 2018 Jul 27];16. Available from: https://www.ncbi.nlm.nih.gov/pmc/articles/PMC5904973/

57. LoopBio. www.loopbio.com. Accessed 17 February 2019.

58. Veedin Rajan VB, et al. Unpublished.

59. Hauenschild C. Der hormonale einfluss des Gehirns auf die sexuelle Entwicklung bei dem polychaeten Platynereis dumerilii. General and Comparative Endocrinology. 1966 Feb 1;6(1):26–73.

60. Payton L, Perrigault M, Hoede C, Massabuau J-C, Sow M, Huvet A, et al. Remodeling of the cycling transcriptome of the oyster Crassostrea gigas by the harmful algae Alexandrium minutum. Scientific Reports. 2017 Jun 14;7(1):3480.

61. Backfisch B, Rajan VBV, Fischer RM, Lohs C, Arboleda E, Tessmar-Raible K, et al. Stable transgenesis in the marine annelid Platynereis dumerilii sheds new light on photoreceptor evolution. PNAS. 2013 Jan 2;110(1):193–8.

62. Reppert SM, Weaver DR. Coordination of circadian timing in mammals. Nature. 2002 Aug 29;418(6901):935–41.

63. Tosini G, Menaker M. Circadian rhythms in cultured mammalian retina. Science. 1996 Apr 19;272(5260):419–21.

64. Balsalobre A, Damiola F, Schibler U. A serum shock induces circadian gene expression in mammalian tissue culture cells. Cell. 1998 Jun 12;93(6):929–37.

65. Dibner C, Schibler U, Albrecht U. The mammalian circadian timing system: organization and coordination of central and peripheral clocks. Annu Rev Physiol. 2010;72:517–49.

66. Schibler U, Ripperger J, Brown SA. Peripheral circadian oscillators in mammals: time and food. J Biol Rhythms. 2003 Jun;18(3):250–60.

67. Fingerman M. Persistent Daily and Tidal Rhythms of Color Change in Callinectes sapidus. Biological Bulletin. 1955;109(2):255–64.

68. Fingerman M, Rao KR, Ring G. Restoration of a Rhythm of Melanophoric Pigment Dispersion in Eyestalkless Fiddler Crabs, Uca Pugilator (Bosc), At a Low Temperature 1). Crustaceana. 1969 Jan 1;17(1):97–105.

69. Fingerman M, Yamamoto Y. Daily Rhythm of Melanophoric Pigment Migration in Eyestalkless Fiddler Crabs, Uca Pugilator (Bosc). Crustaceana. 1967 Jan 1;12(3):303–19.

70. Darnell MZ. Ecological physiology of the circadian pigmentation rhythm in the fiddler crab Uca panacea. Journal of Experimental Marine Biology and Ecology. 2012 Sep 1;426–427:39–47.

71. Stevens M, Rong CP, Todd PA. Colour change and camouflage in the horned ghost crab Ocypode ceratophthalmus. Biological Journal of the Linnean Society. 2013 Jun 1;109(2):257–70.

72. Stevens M. Color Change, Phenotypic Plasticity, and Camouflage. Frontiers in Ecology and Evolution. 2016;4:51.

73. Shahidi R, Williams EA, Conzelmann M, Asadulina A, Verasztó C, Jasek S, et al. A serial multiplex immunogold labeling method for identifying peptidergic neurons in connectomes. eLife. 2015 Dec 15;4:e11147.

74. Nery LE, Da Silva MA, Castrucci AM. Possible role of non-classical chromatophorotropins on the regulation of the crustacean erythrophore. J Exp Zool. 1999 Nov 1;284(6):711–6.

75. Schmid B, Helfrich-Förster C, Yoshii T. A new ImageJ plug-in “ActogramJ” for chronobiological analyses. J Biol Rhythms. 2011 Oct;26(5):464–7.

